# Dissociating proprioceptive deficits in Autism Spectrum Disorders: Intact acuity but impaired sensory integration in postural control

**DOI:** 10.1101/644617

**Authors:** Michail Doumas, Rebekah Knox, Cara O’Brien, Chesney E. Craig

**Affiliations:** School of Psychology, Queen’s University Belfast, Belfast, UK; Research Centre for Musculoskeletal Science and Sports Medicine, Department of Exercise and Sport Science, Manchester Metropolitan University, Crewe, Cheshire, UK

**Keywords:** Proprioception, Postural control, Balance, Autism Spectrum Disorder, Adaptation, Sensory Integration

## Abstract

We investigated the presence of proprioceptive deficits in adults with Autism Spectrum Disorder (ASD), by assessing peripheral proprioceptive information (or proprioceptive acuity) as well as integration of proprioceptive information in the context of postural control. We hypothesized that proprioceptive acuity would be intact but that integration during a postural control task would be impaired. Sixteen adults with ASD and sixteen Neurotypical (NT) adults were screened using an IQ test and the adolescent-adult sensory profile. Proprioceptive acuity was assessed using an ankle Joint Position Sense (JPS) task and integration of proprioceptive information was assessed using a postural adaptation task. This task comprised standing upright, without vision in three phases: standing on a fixed surface for 2 minutes (baseline), followed by standing on a surface tilting in proportion to participants’ body sway, or support-surface sway reference for 3 minutes (adaptation) and finally standing on the restored fixed surface for 3 minutes (reintegration). Results showed no group differences in proprioceptive acuity and in the baseline phase, but greater postural sway during adaptation in individuals with ASD compared with NT controls. Specifically, group differences were not present in the first 30s of adaptation, but emerged after the second window suggesting a deficit in sensory integration of proprioception in adults with ASD. Our results suggest that peripheral proprioceptive information is intact in ASD but neural sensory integration of proprioception is impaired in this group.

---

Autism Spectrum Disorder (ASD) is a life-long developmental disorder primarily diagnosed on the basis of deficits in social interaction and communication as well as restricted and repetitive behaviours (American Psychiatric Association, 2013). In addition, recent diagnostic criteria include atypical sensory processing as one of the characteristic sub-categories of ASD included in DSM-V. These criteria are in line with well-established evidence for impaired sensory function which affects 92% of individuals with ASD (Tomchek & Dunn, 2007) as well as related motor function (Green et al., 2002; Kindregan, Gallagher, & Gormley, 2015; Lim, Partridge, Girdler, & Morris, 2017). The large majority of studies on sensorimotor processing difficulties in ASD have focused on tasks requiring visual processing (for reviews see Black et al., 2017; Gowen & Hamilton, 2013) and visual-auditory integration (for review see Beker, Foxe, & Molholm, 2018), however, the presence of both sensory and motor difficulties in this disorder suggests that proprioception, which is a key sensory channel for movement may also be affected by ASD.

Proprioception refers to sensory information from receptors mainly involved in the senses of limb position and body movement (Proske & Gandevia, 2012). Proprioceptive acuity can be directly assessed using Joint Position Sense (JPS) tasks in which participants are asked to match the position of a limb, either with the other limb (contralateral concurrent matching) or with the same limb a few seconds later (ipsilateral remembered matching)(Goble, Coxon, Wenderoth, Van Impe, & Swinnen, 2009). Given the importance of this sensory channel for movement and daily function it is surprising that only one study has assessed proprioceptive acuity in ASD (Fuentes et al. 2011). This study used an elbow flexion-extension task in adolescents with ASD and Neurotypical (NT) controls. They were asked to either move a line projected in front of them using a joystick to match the position of their unseen arm (passive condition), or to match the position of their unseen arm with the line (active condition). The latter condition was also performed with the index finger. No differences between NT and ASD participants were observed suggesting that primary, peripheral proprioceptive processing (or proprioceptive acuity) is intact in ASD. This result is surprising because people with ASD exhibit general motor deficits (Whyatt & Craig, 2013) but also specific deficits in postural control, especially when proprioceptive information is manipulated alone, or in combination with another sensory channel (Doumas, McKenna, & Murphy, 2016; Minshew, Sung, Jones, & Furman, 2004; Morris et al., 2015). However, Fuentes et al. (2011) only assessed upper limb proprioception, and perhaps there are ASD-related deficits in lower limb proprioception, which is involved in postural control. However, no studies have assessed lower-limb proprioception in ASD.

Postural control relies heavily on proprioception, which has been shown to contribute up to 70% of sensory information in this task (Peterka and Cenciarini, 2006), and on information from visual and vestibular channels. Information from the three channels is integrated using an adaptive sensory integration mechanism with each channel weighted based on its relative reliability (Ernst & Banks, 2002; Peterka, 2002). When environmental conditions change, for example when we step from a hard to a soft surface, the reliability of proprioceptive information for posture decreases. As a result of this change, the weight of the proprioceptive channel decreases and the weights of the two other channels increase to maintain upright stance. Evidence for intact proprioceptive acuity (Fuentes, Mostofsky, & Bastian, 2011) but impaired postural control in ASD (Lim et al., 2017) could suggest that whilst afferent proprioceptive signals may be intact in people with ASD, the way the sensory integration mechanism, integrates or weights this information may be different in ASD compared with NT individuals, and as a result postural sway increases.

Evidence from sensory integration studies supports this idea by showing that individuals with ASD show an over-reliance on proprioception for movement and postural control compared with NT individuals who tend to rely more on vision (Haswell, Izawa, Dowell, Mostofsky, & Shadmehr, 2009; Izawa et al., 2012; Marko et al., 2015; Morris et al., 2015; Sharer, Mostofsky, Pascual-Leone, & Oberman, 2016). For instance, in upper-limb motor learning, individuals with ASD are more sensitive to learning through errors of proprioception than errors of vision (Marko et al., 2015). This over-reliance has also been shown in studies comparing NT and ASD individuals using the Rubber Hand Illusion (Botvinick & Cohen, 1998). Adults with ASD were less likely than NT controls to estimate the location of their actual hand as closer to the rubber hand, rendering their proprioceptive estimates more accurate than NT controls (Paton, Hohwy, & Enticott, 2012). Similarly, children with ASD exhibited delayed susceptibility to this illusion, further indicating inefficient multisensory integration in this population (Cascio, Foss-Feig, Burnette, Heacock, & Cosby, 2012).

ASD-related over-reliance on proprioceptive information has also been shown in postural control tasks (Greffou, 2015; Morris et al., 2015). Oscillations of visual motion were less destabilising for ASD adolescents compared with NT controls suggesting that they relied less on visual cues for balance (Greffou et al., 2012). Similarly, during proprioceptive perturbations using neck vibration ASD adults did not effectively utilise visual information to correct their posture as NT adults did. Instead, they reacted in the same way to the perturbation as when vision was unavailable (Morris et al., 2015). Seen in the context of sensory reweighting models of postural control (Peterka, 2002; Peterka & Loughlin, 2004), these results suggest that, in individuals with ASD, proprioception is relied upon to a greater extent (or up-weighted) and as a result visual and perhaps vestibular information is down-weighted. Consequently, individuals with ASD exhibit a difference in the way they adapt to a new environment requiring proprioception to be down-weighted and vision to be up-weighted. This observation may reflect a fundamental difference in the way proprioception is used during sensory integration in ASD.

The aim of the present study was to identify the origins of ASD-related deficits in the use of use of proprioceptive information. Proprioception was assessed at the peripheral (acuity) and central (sensory integration) levels of processing using a contralateral ankle Joint Position Sense Task and a postural adaptation task respectively. Based on previous evidence (Doumas et al., 2016; Fuentes et al., 2011; Minshew, Sung, Jones, & Furman, 2004a), we predicted that proprioceptive acuity would be intact in individuals with ASD, however when proprioceptive information was manipulated in the context of postural control, we expected that individuals with ASD would be unable to efficiently integrate this information and as a result they would show greater instability compared with NT controls.

Proprioceptive integration was assessed using postural adaptation to an inaccurate proprioceptive environment, tested in three phases: baseline, adaptation and reintegration (Craig & Doumas, 2019; Doumas & Krampe, 2010). In this task, participants stand on a fixed surface with eyes closed, i.e. with only proprioceptive and vestibular information available (baseline, 2 min). Then, inaccurate proprioceptive information is introduced by tilting the support surface in the anterior-posterior direction (toes-down, toes-up) in proportion to the participant’s body sway (or support-surface sway referencing)(Nashner, 1982). In this phase (adaptation, 3 min), with absent vision and inaccurate proprioception only the vestibular system is accurate causing an increase in body sway at a frequency of 0.1Hz (Peterka & Loughlin, 2004). In NT adults, adaptation is reflected in a gradual reduction in postural sway, attributed to down-weighting of inaccurate proprioceptive, and up-weighting of vestibular information (Doumas & Krampe, 2010; Peterka & Loughlin, 2004). Finally, when the fixed surface is restored (re-integration), another period of instability followed by a reduction in postural sway is witnessed, as reweighting must occur in the opposite direction in order to restore the original channel weights. Based on previous studies, we predicted that individuals with ASD would show a greater amount of sway and less efficient adaptation to this environment due to a deficit in down-weighting proprioceptive information.

## Methods

### Participants

Sixteen NT young adults and sixteen young adults with ASD volunteered to participate in this study (see Table 1 for sample characteristics). All participants reported no major neurological or musculoskeletal disorders and no intake of medication that affects postural control (Mamo et al., 2002). Participants with ASD met Diagnostic and Statistical Manual for Mental Disorders- Fourth Edition (DSM-IV, American Psychological Association, 2013) criteria for ASD. Proof of a diagnosis of high-functioning autism was obtained from a General Physician, Clinical Psychiatrist or Clinical Psychologist. Two of the participants with ASD reported diagnoses of co-occurring conditions, one of Attention Deficit Disorder, and one of Attention Deficit Hyperactivity Disorder. Moreover, all participants had full-scale IQ greater than 80 according to the Wechsler Abbreviated Scale of Intelligence (WASI, Wechsler, 1999). ASD severity was assessed using the Social Responsiveness Scale (SRS, Constantino & Gruber, 2005) which was completed by a parent or someone well known to the participant. SRS provides a valid assessment of autism severity as shown by correlation coefficients greater than 0.64 between SRS and the Autism Diagnostic Interview Revised (Hilton et al., 2007). An SRS score of 76 or above is indicative of severe ASD, a score between 60-75 is indicative of moderate severity ASD and a score of 59 or below is within the range of typical development.

**Table 1.** Sample characteristics.

| Characteristic | NT (n=16) | ASD (n=16) | p-value |
| --- | --- | --- | --- |
|  | Mean (SD) | Mean (SD) |  |
| Age | 23.4 (4.3) | 23.6 (5.3) | .81 |
| Sex (female/male) | 5/11 | 6/10 |  |
| Height (cm) | 173.8 (8) | 174.625 (10.8) | .81 |
| Full scale IQ (WASI) | 119.6 (12.4) | 119.4 (12.7) | .97 |
| SP Low Registration | 33.9 (11.5) | 38.5 (10.3) | .04* |
| SP Sensation Seeking | 48.25 (8) | 38.2 (7) | .01* |
| SP Sensory Sensitivity | 34.75 (5.6) | 41.7 (11.5) | .04* |
| SP Sensation Avoiding | 35.1 (8.2) | 45.4 (10.7) | .01* |
| SRS |  | 70 (17.7) |  |

*SP: Sensory Profile. SRS: Social Responsiveness Scale. \*p<.05. Independent samples t-tests were used when normality assumptions were met (Height, SP Sensation Seeking, Sensory Sensitivity and Sensation Avoiding). In comparisons in which normality assumptions were violated (Age, IQ and SP Low Registration) Mann-Whitney tests were used instead.*

Self-reported sensory processing was measured using the adolescent/adult Sensory Profile (Brown & Dunn, 2002), which is a standardised trait measure of sensory processing. The sensory profile is a 60-item self-questionnaire in which an individual answers how he or she generally responds to sensations. Independent samples t-tests revealed group differences in all four categories, with three categories showing greater scores for participants with ASD (SP Low Registration, Sensory Sensitivity and Sensation Avoiding). This is in line with the findings by Kern et al. (2007), who found that individuals with ASD showed greater scores compared with NT individuals. However, one category showed that participants with ASD had lower scores compared with NT participants (Table 1). Participants provided written informed consent and the study was approved by the institution’s Faculty Ethics Committee.

### Apparatus and Tasks

Proprioceptive acuity was measured using an active concurrent contralateral Joint Position Sense task (Fig. 1A). During the task, participants were asked to move their non-dominant foot to match the ankle joint angle of their stationary, dominant foot, and to press a push button confirming foot position when this happened. The JPS task was performed using a custom made device (Fig. 1A) comprised of two thin wooden paddles, attached to linear potentiometers which acted as ankle position transducers. The voltage output signal was converted to angular displacement information with a resolution of 0.0001° (Craig, Goble, & Doumas, 2016). Signals from the potentiometers and the push-button were recorded using a NI-DAQ 6210 card at a frequency of 1000 Hz using custom-written software in MATLAB. Participants were seated in a height-adjustable chair so that their hip and knee angles were at 90° and their feet were strapped to the paddles. The reference (dominant) foot was placed on a fixed height support of either 10° or 15° above horizontal and the reference foot started at 10° below horizontal for all trials. Participants wore a blindfold and headphones during testing and were given a hand-held push button which they were instructed to press when they felt that the matching foot had reached the same level as the reference foot.

**Figure 1.**
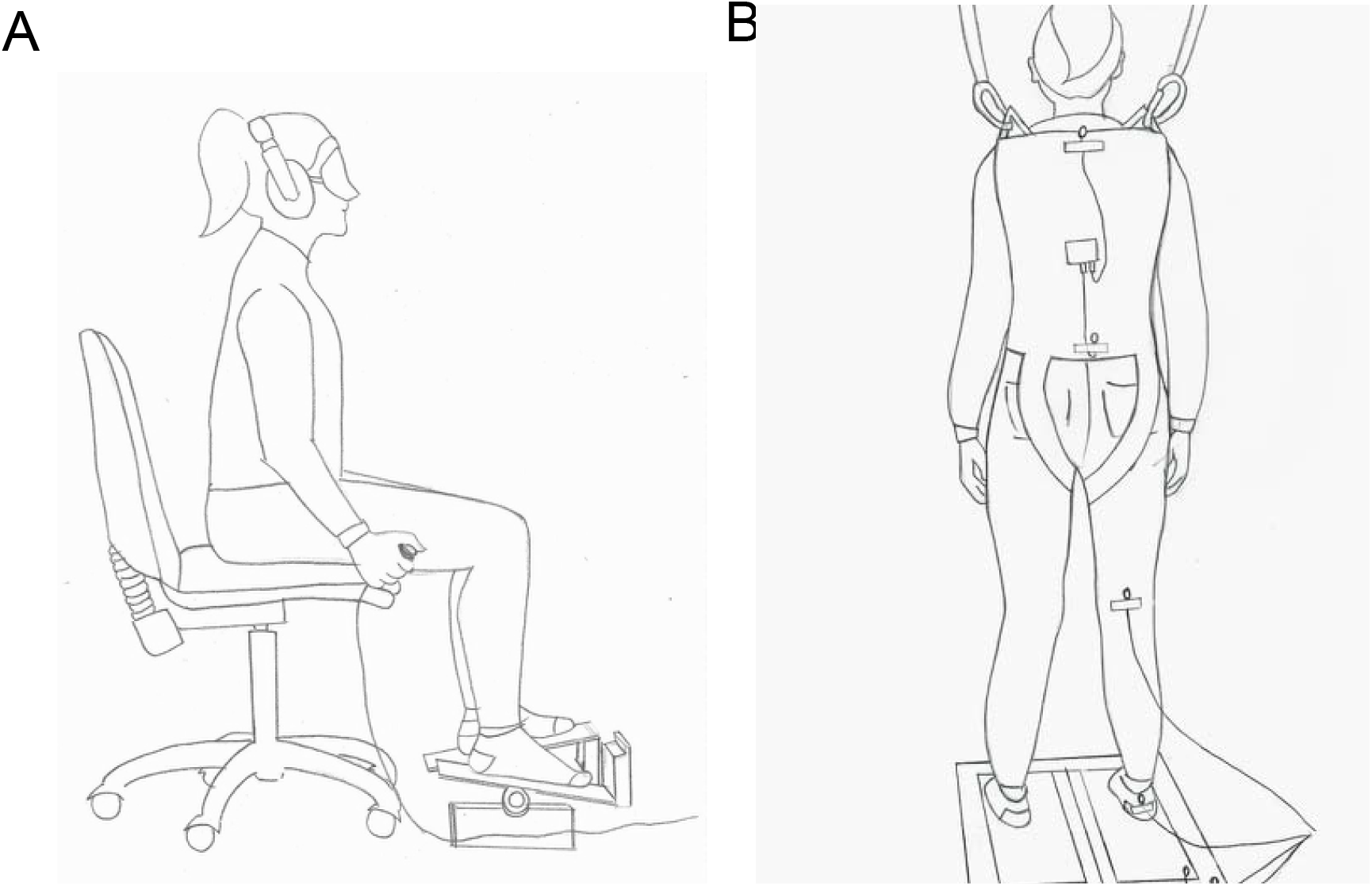
Experimental setups for the two main tasks used in this study. A) Joint Position Matching task and B) Postural control task.

Postural control was assessed using a SMART Balance Master system (NeuroCom International, Inc., Clackamas, OR, USA) comprising an 18”x18” dual force plate and a 3-sided visual surround (Fig. 1B). Stance width was adjusted to each participant’s height in a standardised position according to the system’s manufacturer. Participants wore a harness that was attached to the frame of the Neurocom system in order to prevent a fall, but did not restrict movement, and were instructed to stand upright and to maintain balance on the surface of the force plate. The surface was either fixed or tilted around the ankle joint axis (See Figure 2) in the sagittal plane (Anterior-Posterior direction) in proportion to participants’ expected COM sway angle, also known as support-surface sway reference (Nashner, 1982). COM angles were estimated from the filtered COP-Y trajectory, with a cut-off frequency of 0.5 Hz (Winter, Prince, Frank, Powell, & Zabjek, 1996). A gain setting of 1.6 was used, meaning that body sway of 1° forward resulted in a platform tilt of 1.6 degrees forward, thereby rendering proprioceptive information from the ankle joint inaccurate. This setting was chosen because it has been used previously to demonstrate postural sway differences between ASD and NT adults (Michail Doumas, McKenna, & Murphy, 2016).

**Figure 2.**
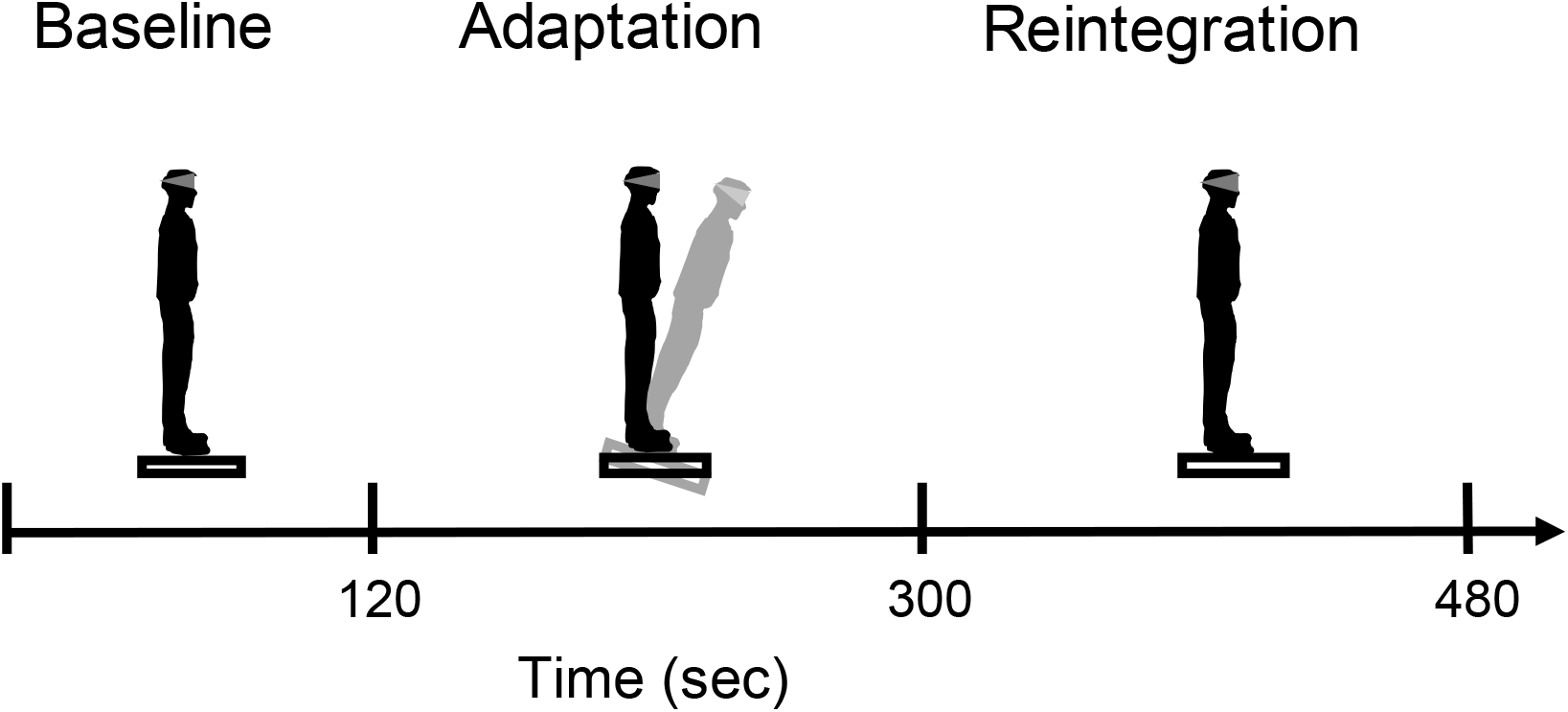
Schematic representation of the postural control task

During sway-referencing, a number of participants in the ASD group displayed loss of stability, which was followed by a small step off the moving surface. When this happened, the trial was interrupted and repeated, and data from this trial were discarded. However, one participant in the ASD group was unable to maintain stability during sway-referencing in repeated trials and was unable to complete the task, therefore this participant’s data were excluded from the sample.

Body kinematics were recorded using a CODA MPX 30 (Charnwood Dynamics Ltd., Leicestershire, UK) three-dimensional (3-D) motion capture system. Infrared light-emitting diodes (ILEDs) were placed at the level of participant’s C7 and L5 vertebrae, on the right knee and ankle, and on the moving and fixed surfaces of the platform.

### Procedure

The experiment took place over 2 sessions, spaced no more than one week apart. The first session took place in the individual’s home or in the laboratory and involved the 4 pre-screening measures: medical information, IQ test (WASI), Sensory Profile and SRS. The second session took place in the laboratory and involved assessment of proprioception (JPS) and balance, which lasted about 1 hour. The JPS task was performed first and consisted of 3 practice trials with the reference foot at an angle of 10° above horizontal (toe-up) which were not included in data analysis, followed by 10 experimental trials with the reference foot at an angle of 15° above horizontal. All trials lasted a maximum of 10 seconds. The start of the trial was signalled by an auditory tone, then participants moved the matching (dominant) foot to what they perceived to be the same position as the reference (non-dominant) foot, and pressed the response button. After the 10s had elapsed a second tone signalled the end of the trial. Between experimental trials, the reference foot was lowered by the experimenter to a height of 10° below horizontal and allowed to rest for 5 seconds before being raised back to the 15° above horizontal height. Participants were then offered a 5 minute break.

The balance task (Figure 2) started with two one-minute practice trials during which the platform was sway-referenced at a gain level of 1.6. The first practice trial was performed with eyes open and in the second participants were asked to wear a blindfold. The practice trial was followed by a single experimental trial which comprised 3 conditions (Figure 2). Participants wore a blindfold and the trial started with standing on a fixed surface for 2 min (Baseline) immediately followed by standing on a sway referenced surface for 3 min (Adaptation) and finally on a fixed surface again for 3min (Reintegration). Participants were instructed before the experiment that they would be warned 10 seconds before the platform began to tilt (start of adaptation) to reduce anticipation and possible anxiety, but would not be told when the platform would stop moving. Participants were not given any information about the manner in which platform movements were generated. Participants were fully debriefed at the end of the study.

### Data Analysis

Proprioceptive acuity was measured as the absolute (absolute angular disparity between the matching and reference feet in a single trial) and variable error (standard deviation of error across all trials of one participant). Gaps (<500 ms) in the motion tracking data from each marker were interpolated using a cubic spline routine in Matlab. Position-time trajectories of motion capture markers were low-pass filtered at 4 Hz using a 5th order Butterworth dual-pass filter. Postural performance was assessed using the lower body angle, or ankle angle in the sagittal plane (AP direction) using kinematic information from the ankle and L5 markers in the y (AP) and z (vertical) dimensions for the 8-minute trial. The ankle angle for the 8-minute trial was segmented into sixteen 30s time windows and Standard Deviation (SD) was calculated in each window. SD has been previously used in studies assessing postural control in ASD (Wang et al., 2016). Spectral properties of the ankle angle waveforms in the first (a1) and last (a6) windows of adaptation were also calculated. This analysis yielded the amplitude-frequency spectrum in bins of 0.033⍰Hz. Amplitude at frequencies up to 0.4⍰Hz (12 frequencies) was calculated as this is the frequency range with the highest amplitude in this task (Doumas & Krampe, 2010; Peterka & Loughlin, 2004). Then, amplitude was averaged in four frequency bins: 0.033–0.1⍰Hz, 0.133–0.2⍰Hz, 0.233– 0.3⍰Hz and 0.333–0.4⍰Hz. We refer to these bins as 0–0.1, 0.1–0.2, 0.2–0.3 and 0.3– 0.4 Hz for simplicity. Data processing was performed using Matlab R2018a (The Mathworks, Natick, MA, USA) and statistical analyses using JASP (Version 0.9.2). Statistical comparisons comprised independent samples t-tests and Mixed Design analyses of Variance. Where sphericity assumptions were violated a Greenhouse-Geisser correction was applied and where normality assumptions were violated non-parametric tests were used. Significance α level was set at 0.05.

## Results

### Proprioceptive acuity

Absolute and variable error results in the JPS task for the two groups are depicted in Figure 3. No group differences were observed, neither for absolute Mann-Whitney U= 172, p=.102, nor for variable error t(30) = 0.83, p=.4, d=.4.

**Figure 3.**
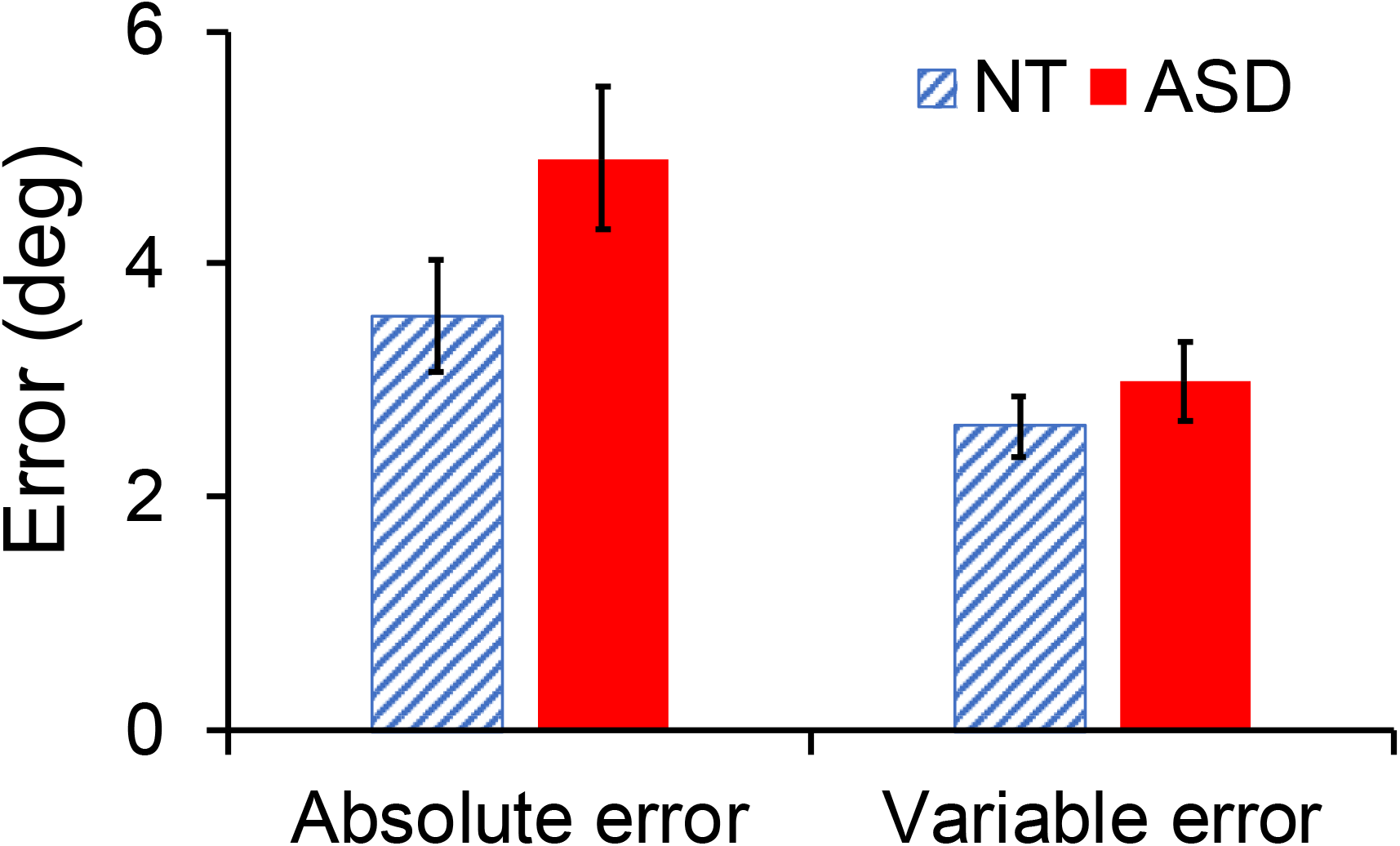
Absolute and variable error results from the JPS task. Error bars depict ± 1 standard error of the mean.

### Postural control (Ankle Angle SD)

Raw ankle angle data from a representative NT and a representative ASD participant as a function of time are depicted in Figure 4. Visual inspection of the figure shows greater sway amplitude for the ASD participant in the adaptation phase compared with the NT participant. Average results for ankle angle SD in 30s windows for the three phases are depicted in Figure 5. Below we report group and window comparisons of ankle angle SD for each of the three phases separately.

**Figure 4.**
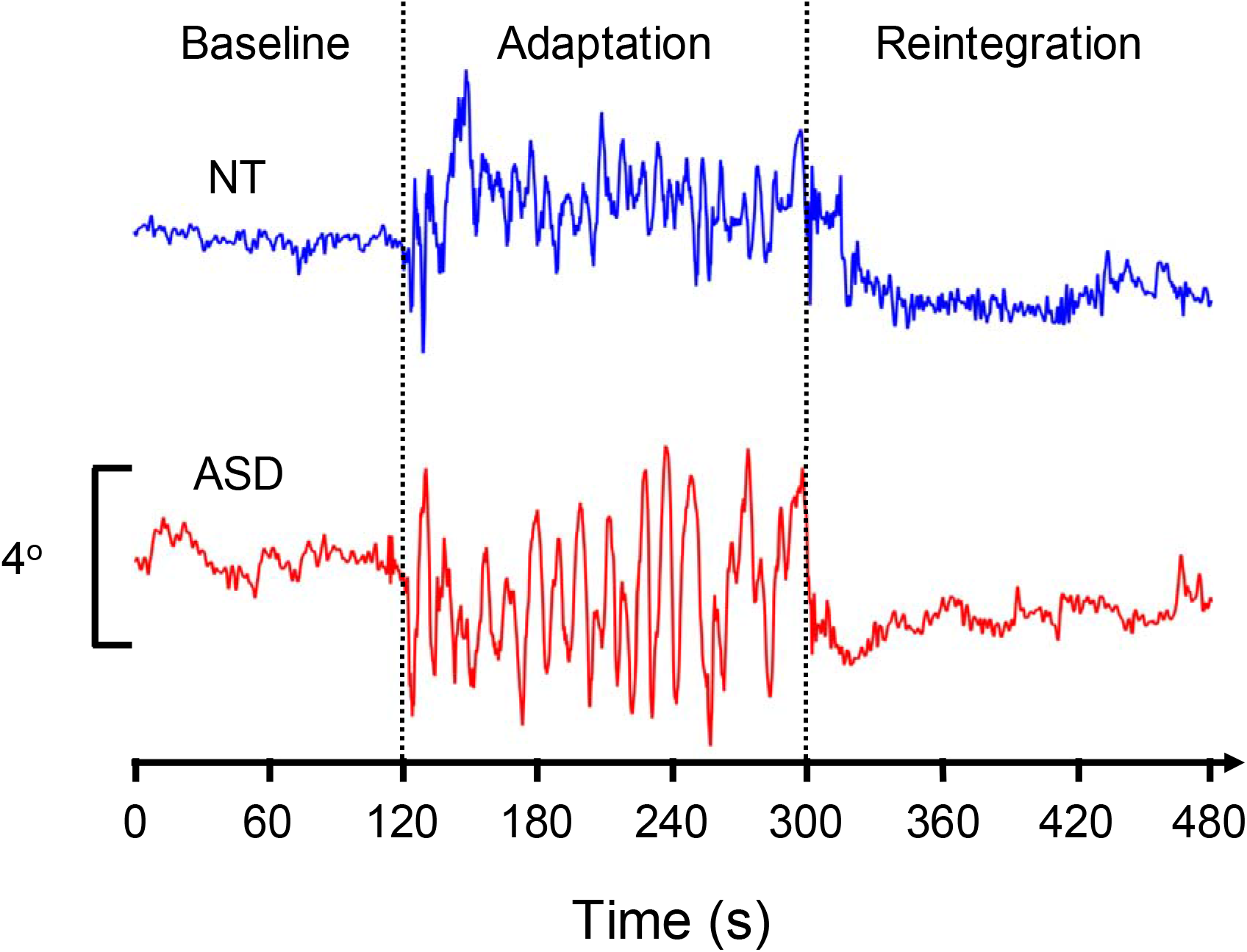
Ankle angle trajectories over time for the three phases of the postural control task in one representative NT participant (top) and one representative participant with ASD (bottom).

**Figure 5:**
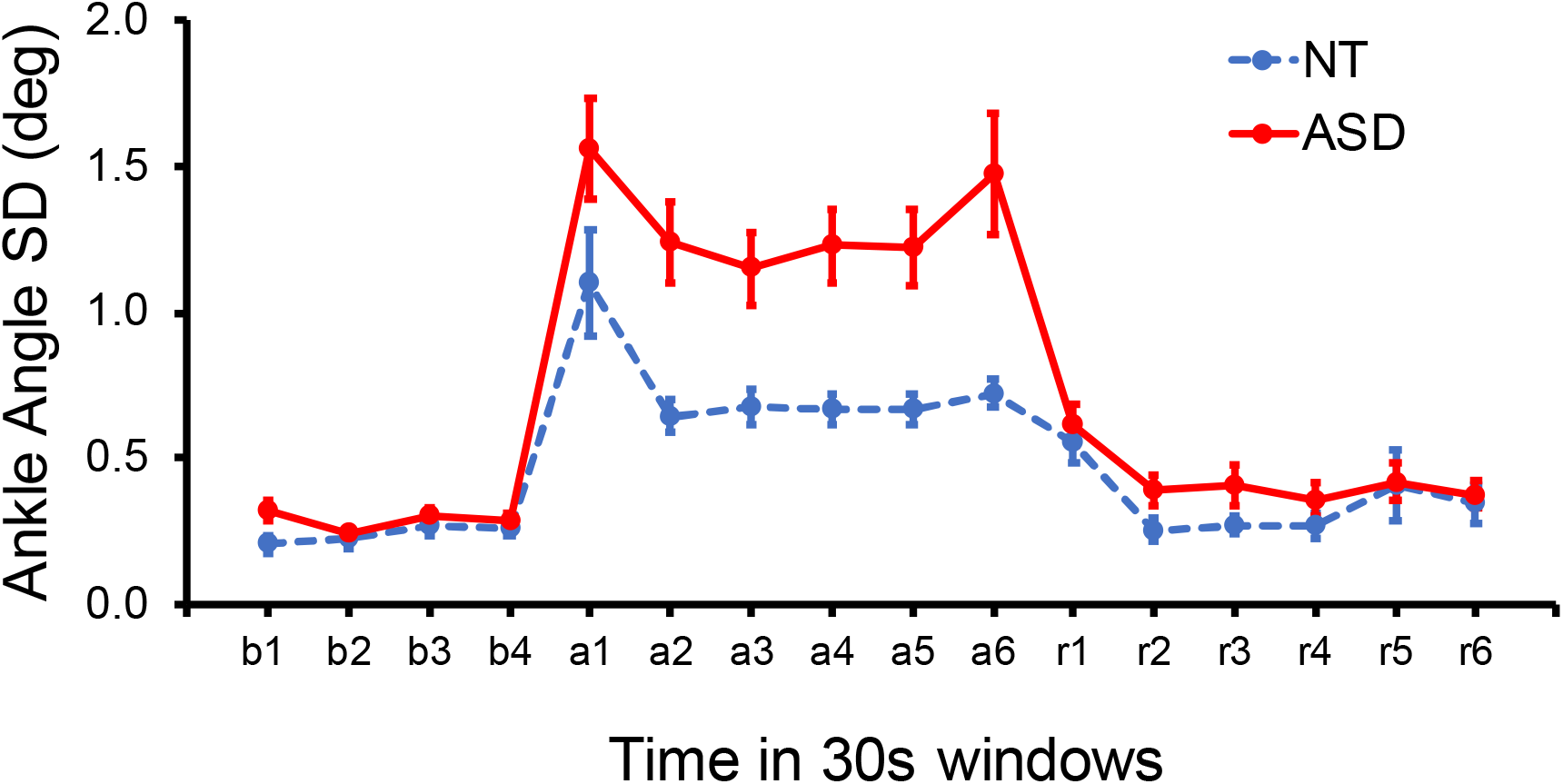
Ankle angle SD in successive 30s windows over the three phases: Baseline (b1-b4), Adaptation (a1-a6) and Reintegration (r1-r6). Error bars depict ± 1 standard error of the mean.

### Baseline

Ankle angle SD at baseline (Fig. 5, windows b1-b4) was analysed using a 2 by 4 Mixed design ANOVA with group (NT and ASD) as between- and window (1-4) as within-subjects factors. No significant main effects or interaction were shown.

### Adaptation

When sway referencing was introduced, a 2 by 2 Mixed design ANOVA contrasting ankle angle SD in the two groups between the last window of baseline with the first window of adaptation (b4 vs. a1) was performed. Ankle angle SD showed a large increase when sway reference was introduced reflected in a main effect of window F(1,30) = 66.51, P<.001, η^2^ =.67 but there was no group difference during this transition as shown by the lack of a main effect of group and a window by group interaction.

Throughout the adaptation phase (Fig. 5, windows a1-a6), ankle angle SD was greater in the ASD compared with the NT group (M_NT_=0.7°, SD_NT_=0.3°, M_ASD_=1.3°, SD_ASD_=0.4°) as shown by a main effect of Group F(1,30) = 21.56, P<.001, η^2^ =.418. Pair-wise comparisons in each window corrected for multiple comparisons (6 comparisons, adjusted α=0.0083) showed that there were no differences in window a1 t(30) = 1.83, P=.077, but in all other windows angle SD was greater in the ASD group (t(16.7-21.2)=3.5-4.1,all P<.0083). Overall, angle SD decreased over time as shown by a main effect of window F(3,88.8) = 5.24, P<.001, η^2^ =.15. No window by group interaction was shown.

### Reintegration

To assess the presence of an aftereffect when the fixed platform was restored we performed pair-wise t-tests contrasting the mean ankle angle SD at baseline (windows b1-b4, M_NT_=.220°, SD_NT_=0.1°, M_ASD_=.289°, SD_ASD_=0.07°) with each of the reintegration windows, separately for the two groups. Results showed that in both groups a significant aftereffect was shown in window r1 NT: t(15) = 4.63, p<.001, d=1.16; ASD: t(15) = 5.34, p<.001, d=1.33; with smaller effects also shown in the NT group on window r6 t(15) = 2.15, p=.048, d=.51 and in the ASD group on window r5 t(15) = 2.23, p=.041, d=.56; Furthermore, ankle angle SD during reintegration (Fig. 5, windows r1-r6) was analysed using the same ANOVA as for baseline and adaptation. Results showed a reduction in angle SD over time F(2.4,72.7) = 9.22, P<.001, η^2^ =.23 but no group effect or group by window interaction.

### Spectral analysis

Statistical analyses of spectral amplitude of ankle angle trajectories (Fig. 6) in the dominant frequencies in this task (0-0.4Hz) were performed using a mixed design ANOVA with group as between- and window (a1 and a6) and frequency band (0-0.1, 0.1-0.2, 0.2-0.3 and 0.3-0-4) as within-subjects factors. Participants with ASD showed greater amplitude, especially in low frequencies, as shown by a frequency by group interaction F(1.3,38.17) = 11.6, P<.001, η^2^ =.09 and main effects of frequency F(1.3,38.17) = 93.27, P<.001, η^2^ =.7 and group F(1,30) = 7.62, P=.01, η^2^ =.2. Pair-wise comparisons of the two groups in all frequency bands corrected for multiple comparisons (8 comparisons, adjusted α=0.00625) showed that group differences were only observed in the last adaptation window (a6), in the lowest (0-0.1) frequency band t(17.14) =3.32, p=0.004, d=1.17.

**Figure 6:**
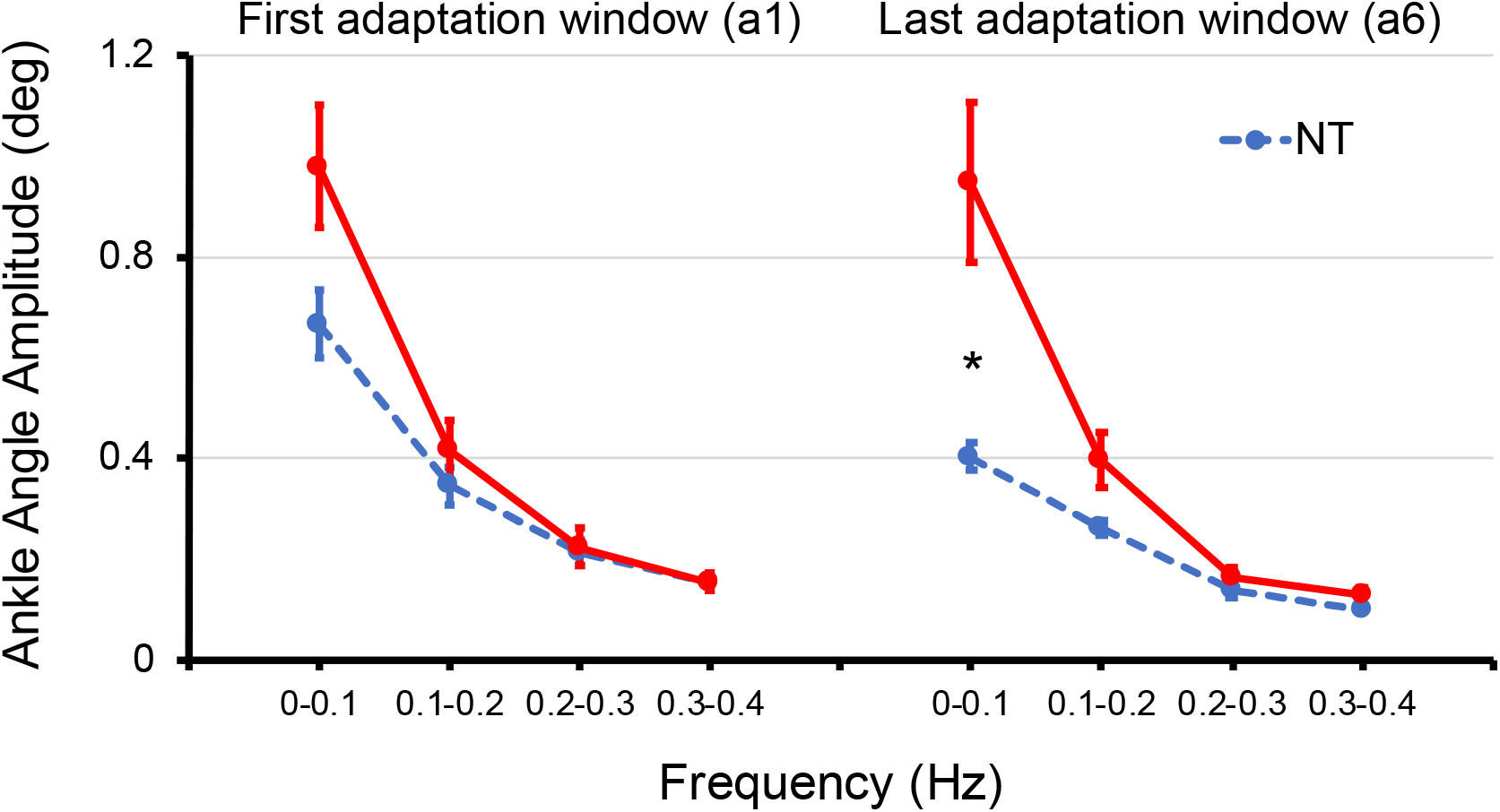
Spectral properties of ankle angle trajectories the first (left) and the last (right) 30s window of adaptation in the two groups. Error bars depict ± 1 standard error of the mean. *Significant group differences (p<.05).

## Discussion

The aim of this study was to assess the dissociation between intact proprioceptive acuity and impaired proprioceptive processing in postural control in individuals with ASD. Results showed that there were no differences between NT and ASD individuals in proprioceptive acuity. Furthermore, there were no group differences in baseline postural control with eyes closed or when the inaccurate proprioceptive environment was initially introduced. However, after the first 30s of adaptation, NT participants showed a marked reduction in postural sway reflecting optimal adaptation, whereas sway levels in participants with ASD remained high. This group difference was evident both in postural sway overall (ankle angle SD) and in sway at low frequencies (0-0.2Hz), which is the frequency targeted by the sway referencing manipulation (Clark & Riley, 2007; Doumas & Krampe, 2010; Doumas, Valkanidis, & Hatzitaki, 2019; Peterka & Loughlin, 2004). Overall, our hypothesis for intact proprioceptive acuity but impaired proprioceptive integration in ASD was supported.

Our finding that ASD individuals displayed greater sway than NT individuals in response to challenging sensory conditions is strongly supported in the literature (Bucci et al., 2017;Doumas et al., 2016; Molloy et al., 2003; Price et al., 2012; Stins et al., 2015; Travers et al., 2018). Moreover, the demonstration of a significant group difference after the first 30s of the adaptation phase is particularly striking, as it shows that participants with ASD could not adapt to the proprioceptive manipulation over a 3-minute period. This could not be explained by a general postural deficit, as there were no group differences at baseline nor during the reintegration phase. Furthermore, our previous research has shown that this may be unique to the proprioceptive channel, as visual surround sway-referencing did not show this effect (Doumas, McKenna, & Murphy, 2016). This suggests that the current results do not merely reflect an inefficient sensory integration process, but a more specific deficit in the ability to down-weight unreliable proprioceptive information. This inability to adapt to a proprioceptive manipulation supports previous evidence that individuals with ASD are overly reliant on proprioception for movement and postural control (Haswell, Izawa, Dowell, Mostofsky, & Shadmehr, 2009; Izawa et al., 2012; Marko et al., 2015; Morris et al., 2015; Sharer, Mostofsky, Pascual-Leone, & Oberman, 2016).

Despite the confirmation of our main hypothesis, alternative explanations for adaptation deficits in the ASD group cannot be discounted. Firstly, ASD-related deficits in the vestibular system could contribute these findings. This is because during adaptation to an environment with inaccurate proprioceptive information and no vision (adaptation phase) the vestibular channel is the only reliable source of sensory information (e.g. Peterka et al., 2002). Unfortunately, evidence regarding the integrity of the vestibular system in ASD is limited and inconsistent (e.g. Carson et al., 2017). Peripheral vestibular function has been investigated in ASD through examination of the vestibular ocular reflex (VOR; Cullen, 2012). Early experiments reported abnormal rotational VOR (rVOR) in children with ASD compared to NT controls (Ornitz, 1970; Ornitz et al., 1974; Ornitz et al., 1985; Ritvo et al., 1969) and similar findings are reflected in the most recent study on the topic (Carson et al., 2017). However, in two other recent studies, no differences rVOR were detected in large samples of children or adults with high-functioning ASD compared to NT controls (Goldberg et al., 2000; Furman et al., 2015). Thus, there is no clear consensus on the intactness of peripheral vestibular processing in ASD. Moreover, to our knowledge, postural control during vestibular manipulations has not yet been measured in this population. Therefore, future research into both peripheral vestibular processing and vestibular processing during balance in ASD individuals is warranted to probe this explanation.

Another explanation may relate to the general sensory integration demands of a task required to demonstrate postural control differences between ASD and NT individuals. It is possible that the addition of a proprioceptive manipulation on top of already absent vision raised general sensory integration requirements above the threshold required to reveal ASD-related deficits in adaptation. Future studies should investigate this possibility by measuring adaptation to manipulations of vision, vestibular sense and proprioception both separately and in pairs. This will confirm whether only manipulations involving proprioception produce reveal ASD increases in sway, or whether perhaps paired manipulations, in general, increase sensory integration demands enough to prevent sway adaptation in ASD. However, based on currently available evidence from postural control (Doumas et al., 2016; Minshew et al., 2004; Morris et al., 2015) and other motor tasks (Haswell et al., 2009; Izawa et al., 2012; Marko et al., 2015; Sharer et al., 2016) a specific sensory integration deficit in the form of a bias for proprioceptive processing remains the most convincing proposal.

The bias in processing proprioceptive input observed here, even when it is maladaptive, could suggest a neural deficit whereby the sensory reweighting mechanism may be deficient at inhibiting variable proprioceptive inputs. This is interesting considering that increasing evidence supports an excitatory/inhibitory (E/I) imbalance in neural networks as the underlying cause of many ASD symptoms, including social problems, stereotyped behaviours and perceptual and motor deficits (Nelson & Valakh, 2015). Similarly, regional cortical E/I imbalances are proposed as an underlying pathology in attention-deficit hyperactivity disorder (ADHD; Krause, Márquez-Ruiz, & Cohen Kadosh, 2013), in which deficits in sensory reweighting are also presented (Bucci et al., 2017). Whilst the neural pathology for sensory reweighting deficits in these neurodevelopmental disorders is currently unknown, future research could investigate this using non-invasive neuroimaging techniques, such as, electroencephalography (EEG) or functional near-infrared spectroscopy (fNIRS) during sensory manipulations.

### Limitations

Our study had a number of limitations. Firstly the highly controlled nature of the proprioceptive manipulation may not accurately reflect typical proprioceptive challenges encountered outside the laboratory, such as standing on a moving vehicle. Secondly, we did not measure developmental trajectory and only included individuals with an IQ of 80 or above. Our findings can therefore only be applied to the population of adolescents and young adults with high-functioning ASD. Future studies using more ecologically valid tasks with a more representative ASD sample will be required to fully investigate this finding.

### Future directions

Future interventions should aim to address overreliance on proprioception and relative under-reliance on vision in ASD individuals. One option could be to train individuals with ASD to better-utilise visual cues to maintain balance. For instance, Somogyi et al. (2016) found that providing visual representation of real-time sway information to children with ASD led to significant improvements in postural control. Another, recent study also trained ASD and NT children over 6 weeks using a visual biofeedback-based video game, finding improvements in postural stability both within and outside of the training context (Travers et al., 2018). These promising results suggest that, through training, children with ASD may be able to effectively use visual feedback to improve postural control.

In conclusion, our study shows that individuals with ASD weight proprioceptive information for postural control differently to NT individuals, despite displaying normal lower-limb proprioceptive acuity. Our findings suggest that individuals with ASD are unable to down-weight the proprioceptive channel, resulting in an inability to adapt to challenging proprioceptive manipulations. Future research addressing limitations and alternative explanations described here will inform future postural control interventions in this population.

